# An equation-free method reveals the ecological interaction networks within complex microbial ecosystems

**DOI:** 10.1101/080697

**Authors:** Kenta Suzuki, Katsuhiko Yoshida, Yumiko Nakanishi, Shinji Fukuda

## Abstract

1. Mapping the network of ecological interactions is key to understanding the composition, stability, function and dynamics of microbial communities. In recent years various approaches have been used to reveal microbial interaction networks from metagenomic sequencing data, such as time-series analysis, machine learning and statistical techniques. Despite these efforts it is still not possible to capture details of the ecological interactions behind complex microbial dynamics.
2. We developed the sparse S-map method (SSM), which generates a sparse interaction network from a multivariate ecological time-series without presuming any mathematical formulation for the underlying microbial processes. The advantage of the SSM over alternative methodologies is that it fully utilizes the observed data using a framework of empirical dynamic modelling. This makes the SSM robust to non-equilibrium dynamics and underlying complexity (nonlinearity) in microbial processes.
3. We showed that an increase in dataset size or a decrease in observational error improved the accuracy of SSM whereas, the accuracy of a comparative equation-based method was almost unchanged for both cases and equivalent to the SSM at best. Hence, the SSM outperformed a comparative equation-based method when datasets were large and the magnitude of observational errors were small. The results were robust to the magnitude of process noise and the functional forms of inter-specific interactions that we tested. We applied the method to a microbiome data of six mice and found that there were different microbial interaction regimes between young to middle age (4-40 week-old) and middle to old age (36-72 week-old) mice.
4. The complexity of microbial relationships impedes detailed equation-based modeling. Our method provides a powerful alternative framework to infer ecological interaction networks of microbial communities in various environments and will be improved by further developments in metagenomics sequencing technologies leading to increased dataset size and improved accuracy and precision.

## Introduction

Microbial communities contribute to the evolutionary fitness of other living organisms by inhabiting their bodies (Mueller and Sachs 2015) and surroundings (Chaparro et al. 2012; Heederik and von Mutius 2012; Panke-Buisse et al. 2015). For example, the gut microbiota assists host metabolism (Tremaroli and Bäckhed 2012; Sommer and Bäckhed 2013) and provides defense against pathogens (Kamada et al. 2013). This understanding has motivated the development of microbial medicinal interventions that attempt to treat various disorders through the manipulation and control of microbial communities (Borody and Khoruts 2012). Furthermore, an emergent property of the microbial community is the potential contribution to environmental remediation through the degradation of pollutants (Swenson et al. 2000; Iranzo et al. 2001). The composition, stability, function and dynamics of a microbial community provides the mechanistic basis for these microbial treatments, and closer ties between these ecosystem properties and ecological interaction networks (interaction webs) have been revealed (Tylianakis et al. 2010). Hence, an understanding of ecological interaction networks is crucial for both human health and environmental sustainability. While it is difficult to study complex microbial interactions using traditional laboratory cultivation approaches, recent developments in next generation sequencing technology and high performance computing environments have enabled various approaches for revealing ecological interaction networks, ranging from time-series analysis, machine learning and statistical techniques (Faust and Raes 2012; Bucci and Xavier 2014; Faust et al. 2015; Vacher et al. 2016). However, there are currently no sufficiently effective methods for capturing details of ecological interactions within microbial communities, which frequently exhibit complex dynamics (Caporaso et al. 2011; Dethlefsen and Relman 2011; Pepper and Rosenfeld 2012; Relman 2012; Ravel et al. 2013; Gerber 2014).

An ecological interaction network is defined as a directed network that describes interactions between organisms, such as mutualisms, competition and antagonistic (predator-prey) interactions (Morin 2011; Faust and Raes 2012). An ecological interaction network is generally described as a pairwise interaction matrix whose elements take zero, positive or negative values with regard to the effect of one species on the other. Here, we summarize the ties between ecological interaction networks and other ecosystem properties in three main points. First, the stability of an ecological system relies on its ecological interaction network, as is known from the seminal work of May (1974) that formulated how the stability of an ecological system may depend on the density and strength of its ecological interactions. In microbial communities in particular, mutualistic interactions may have a disruptive effect on community stability (Coyte et al. 2015). Second, there are extensive studies suggesting that an interaction network is crucial to the development and maintenance of microbial ecosystem functions (reviewed by Vacher et al. 2016). For example, findings from a type of artificial selection experiment led Blouin et al. (2015) to suggest that reducing interaction richness is crucial to developing and maintaining microbial ecosystem function in terms of low CO_2_ emission. Third, an ecological interaction network of a real ecological system can be used to test the validity of network based metrics for keystone species, i.e. species having significant effects on the stability and/or function of ecological systems out of proportion to their abundance (Paine 1969; Power et al. 1996; Jordan 2009). For example, a “keystone index” identifies keystone species based on their topological position within an interaction network (Jordan 1999), and is a pioneering theoretical development underpinning recent microbial community studies (Berry and Widder 2014; Toju 2016).

An ecological *interaction* network is different from an ecological *association* network such as a correlation network (Friedman and Alm 2012) or co-occurrence network (Faust et al. 2012). Although correlations between time-series data are often used as a proxy for interactions between species, this is not a reliable method even if a strong correlation exists between two species (Fisher and Mehta 2014). A co-occurrence network only implies the presence of underlying ecological interactions, whereas it provides significant information regarding associations between microbial species (Vacher et al. 2016). As an alternative approach, algorithms have been developed to infer ecological interaction networks directly from microbial time-series (Jiang et al. 2013; Fisher and Mehta 2014; Bucci et al. 2016). However, these algorithms may not be applicable to the complex dynamics of microbial communities, which require the following algorithm properties. First, a time-series demonstrating non-equilibrium dynamics must be available because such dynamics are common in microbial interaction networks. There are many reasons for this, such as species interactions, environmental fluctuations, experimental perturbations, invasions and aging, and understanding the dynamics resulting from these processes is clearly an important goal. Second, a method that can capture microbial relationships without any presumption regarding their mathematical formulation (in other words, an equation free modelling approach) is desirable. As claimed for ecological systems in general (Deyle et al. 2016), ecological interactions are often nonlinear, such that the effect of species X on Y is not simply proportional to the abundance of Y, and attempting to formulate all these relationships into mathematical functions is not realistic (DeAngelis and Yurek 2015; Bashan et al. 2016). This fact will reduce the reliability of approaches that assume a *priori* any underlying equation. Overcoming these obstacles will widen the applicability of network inference methods without reducing reliability, and will increase our understanding of microbial communities.

We developed an algorithm, the Sparse S-Map method (SSM; Fig 1, Materials and Methods), that satisfies the above requirements. This algorithm generates a sparse interaction network based on the predictability of an ecological time-series without assuming any particular underlying equation. Using simulated multispecies population dynamics, we compared the performance of the SSM to a comparable equation-based method (LIMITS; Fisher and Mehta 2014) for different dataset sizes, magnitudes of process noise and observational errors, and functional forms of species interactions to highlight the differences between equation-free and equation-based methods. We then applied the SSM to a time-series of gut-microbiota taken from the faeces of six mice over 72 weeks. To harness data limitations (18 time points per mouse), we pooled data from five mice for the predication dataset and used it to infer a network that best explained the dynamics of the remaining mouse.

**Figure 1.**
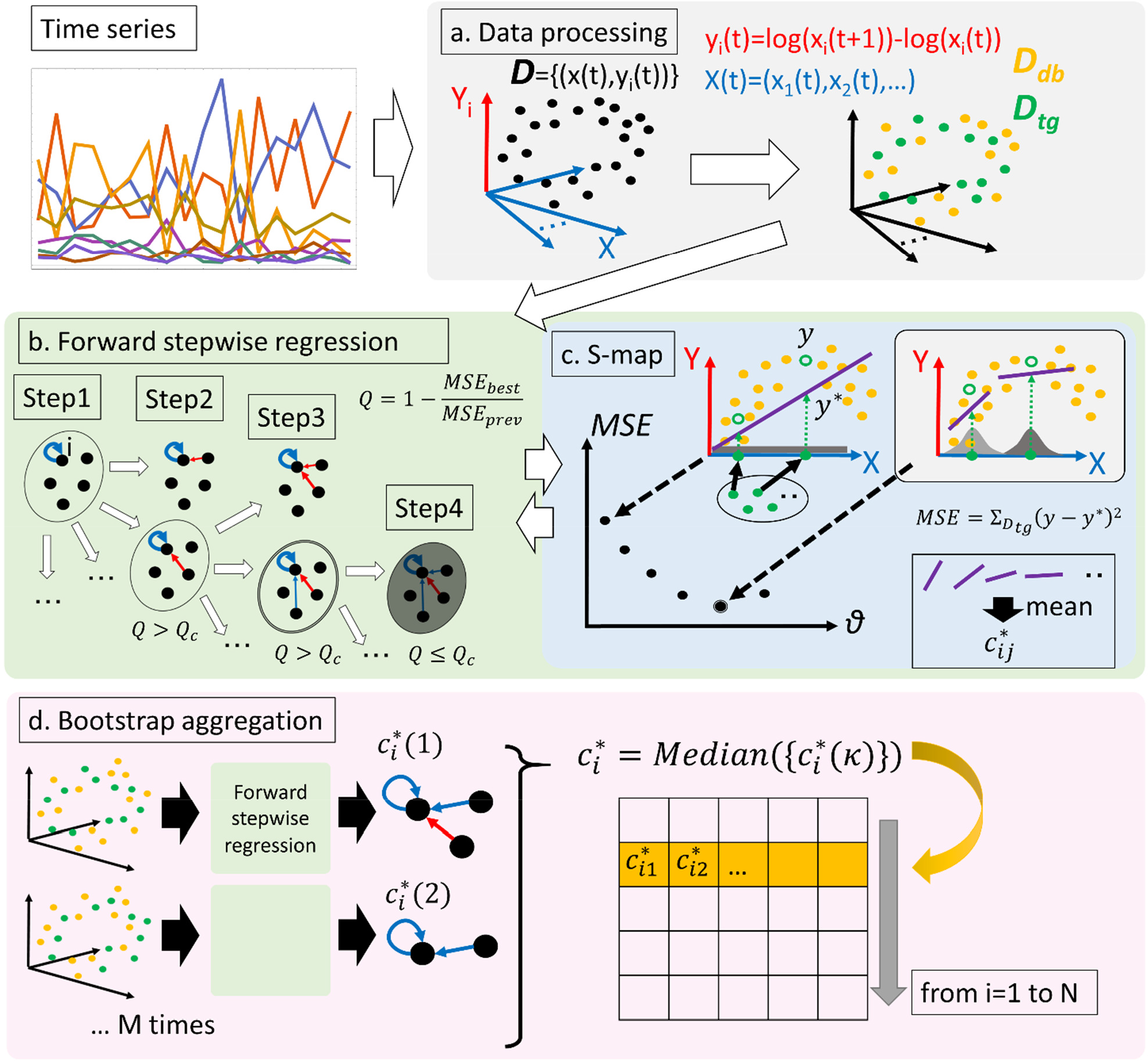
Overview of the sparse S-map method (SSM).

## Materials and Methods

### Sparse S-map method

#### Over view of the Sparse S-map method

The sparse S-map method (SSM) is an algorithm that executes a forward stepwise regression scheme with bootstrap aggregation (“bagging”) to calibrate species’ interaction topology (i.e. with which species a focal species interacts) for S-map (Dixon et al. 1999, Sugihara et al. 1996, Sugihara 1994). In other words, the SSM is a data-oriented equation-free modelling approach (Empirical Dynamic Modelling, EDM; Deyle et al. 2013; Ye et al. 2015; Deyle et al. 2016; Ye and Sugihara 2016) for multispecies ecological dynamics whose interaction topology is unknown. The SSM is essentially a non-parametric method that does not require any additional effort to adjust parameters for the given data. Furthermore, owing to the forward stepwise scheme, it is applicable to both absolute and relative abundance data without special treatment (Fisher and Mehta 2014). Fisher and Mehta (2014) have already applied the forward stepwise scheme with bagging to a linear regression model and thus developed an algorithm inferring the sparse interaction matrix (LIMITS). However, because of the use of a linear regression model, the authors limited the applicability of LIMITS to systems whose dynamics are close to equilibrium.

The SSM is concisely explained as follows. First, in the data processing process, time-series data is processed to a set of pairs of multivariate explanatory variables and a response variable (*D*; Fig. 1a). Also in this step, these pairs are separated into *D_db_* and *D_tg_*, a training set and a testing set. Second, in a forward stepwise procedure, explanatory variables used for the regression are added (Fig. 1b). In this step, S-map is used as the regression method (Fig. 1c). At the end of the forward regression procedure, interaction strengths are obtained as the mean of coefficients from the regression function. Then, in the bootstrap aggregation procedure, the first and second procedures are repeated with randomly generated *D_db_* and *D_tg_*, and the median of the interactions strengths are taken as the final result (Fig. 1d).

#### Data processing

We assume that time-series 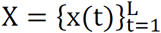 is an array of vectors 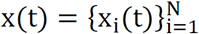 where t = 1, …, L indicates data points with a constant interval (say 1 day), i = 1, …, N indicates species (OTUs) and 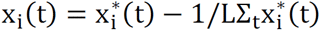 is the abundance of species adjusted to a mean of zero, where 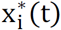 is the abundance. If 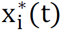 is the relative abundance, then Σ_i_x_i_(t) = 0. However, we do not specify whether x_i_(t) is relative abundance or absolute abundance because our method is applicable to both cases. For convenience, we assume that 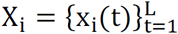 is the time series of species i. We also define a time series 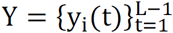, with 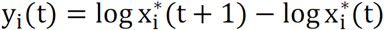, to apply gradient matching (Ellner et al. 2002), which assumes y_i_(t) as the response variable and x(t) as the explanatory variable. In the regression processes, the explanatory variables and the response variable having the same time index is treated as a pair (x(t),y_i_(t)). We refer to the set of these pairs 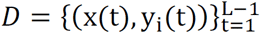 as “data”. Please note that while y_i_(t) is a scalar, x(t) is a vector with N elements.

#### Bootstrap aggregation

Because the forward stepwise regression explained below is known to be unstable, we used a bootstrap aggregation method to obtain a stable result. To apply the bagging procedure, half of the pairs in *D* are randomly sampled to make a “database” *D_db_* and the rest of the points are a “target” *D_tg_*. We use 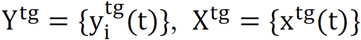 and 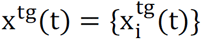 when we need to specifically indicate the points in D_tg_. The process of forward stepwise regression is repeated Z times with different partitioning. Here, we set Z = 100.

#### S-map

S-map is a locally weighted linear regression model used for the mechanistic prediction of complex ecological dynamics (Deyle et al. 2016). S-maps approximate the best local linear model by giving greater weight to points that are close to the current ecosystem state (Fig. S1a). It is applicable to complex ecological dynamics without any limitation in the dynamic property of given data and requires no special effort in formulating the underlying species relationships into mathematical functions. However, so far it has only been applicable to ecological systems with a small number of species whose interaction topology is already known. By applying a forward stepwise regression with bootstrap aggregation, the SSM realizes the appropriate selection of the interaction topology for S-map so that S-map becomes most relevant for explaining a given set of data points. Hence, in the SSM, S-map is applied to a subset of variables (e.g., Fig S1b) in a stepwise manner. The ability of S-map to describe non-linear species relationships makes the selection of interacting species reliable.

##### Algorithm 1: S-map

1. Initiate θ = 0.

2. Select a pair 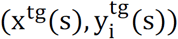 from *D_tg_*.

3. Calculate weight vector by,

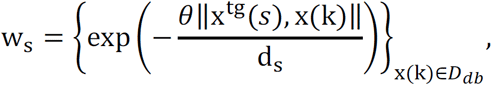
where, 
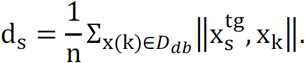

Here, ∥∙,∙∥ denotes the Euclidian distance between two vectors and n = |*D_db_*| is the number of elements in *D_db_*.

4. Generate a weighted design matrix as,

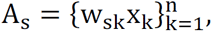
where w_sk_ is the kth element of w_*s*_. Here, 1 is added as the N + 1th element of x_k_s to include the constant term for regression.

Similarly, generate a vector of weighted response variable as,

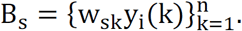

5. Solve a linear equation

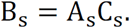
as, 
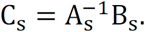
Here, 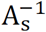 is the pseudoinverse of A_s_.

6. Prediction for 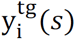 is obtained as,

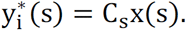

Here again 1 is included as the N + 1th element of x(s).

7. Iterate 2-6 until all pairs in D_tg_ is selected. Then, calculate

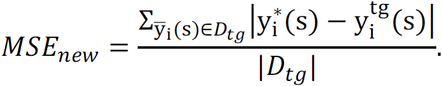

8. If θ = 0, or θ ≠ 0 and *MSE_new_* < *MSE* inclement θ by dθ, set *MSE* = *MSE_new_* and back to 2, else return *MSE*.

The ith column of the inferred interaction matrix, 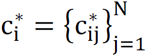 is obtained by simply assuming that

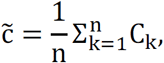
and set 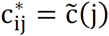 for j ∈ I_active_ and 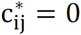 otherwise. Here, 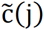 represents the element of 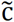 corresponding to the jth species in I_active_. As explained in the next section, I_active_ is the set of the index of species whose interaction to species i is active (thus, non-zero).

The parameter θ tunes how strongly the regression is localized to the region of state space around each 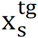. Note that if θ = 0, the S-map model reduces to a vector auto-regression (VAR) model. Thus, S-maps include linear VAR models as a special case. More importantly, this also means that the SSM includes LIMITS as a special case. For θ > 0, the elements of w_s_ can vary with the location in the state space in which (x, y_i_) is plotted, and with increasing θ they can vary more strongly for different xtg. If θ is too small, the coefficients will underestimate the true variability in interaction strength. However, with larger θ the regression hinges on only the most proximal points on the manifold and will therefore be more sensitive to observation error. Here, we selected θ that minimizes *MSE* by incrementing θ from zero by dθ = 0.2 steps because as a function of θ, *MSE* generally has a global minimum not very distant from zero (say, θ < 10). It is the simplest procedure for the minimization of *MSE* adopted for explanation, and would be replaced by a more sophisticated method. For instance, an algorithm that initially searches for the best θ in a broad range with a low resolution (e.g., θ = {0,1, …,8}) and then increases the resolution with a stepwise narrowing of the search range may work as well. θ is used as an indicator for the degree of non-linearity of dynamics (Sugihara 1994; Sugihara et al. 1996). Here, we also used θ as a proxy for non-equilibrium of time-series data (see supplement information).

#### Forward stepwise regression

The use of forward stepwise regression was motivated by two reasons (Fisher and Mehta 2014). The first reason is that the forward stepwise selection can distinguish between the presence and absence of species interactions and include interactions only when it improves the predictive power of the model. This makes inferred interaction networks sparse and easily interpretable. The second reason is that modern metagenomic techniques can only measure the relative abundances of microbes, not their absolute abundances. Hence, the design matrix for the linear regression becomes singular, and there exists no unique solution to the ordinary least squares problem. In the forward stepwise procedure interactions and species are added sequentially to the regression as long as they improve the predictive power of the model. Because the design matrix now only contains a sub-set of all possible species, it is never singular and the linear regression problem is well-defined. Below, we describe the forward stepwise regression including bootstrap aggregation. Since all of the regressions are performed independently for each species, we described the algorithm for a species, i.e. inferring a row of the interaction matrix (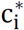). The full interaction matrix is obtained by repeating the procedure from i = 1 to N.

##### Algorithm 2: Forward stepwise regression with bootstrap aggregation

1. Set the target species i.

2. Initiate index z = 1.

3. Sample half of the pairs in *D* to make *D*_db_, and set the rest as *D*_tg_.

4. The set of the index of explanatory variables (species) that have active interactions to species i is initialized to I_active_ = {i} because the presence of intra-specific interaction is natural, and the interaction with the rest of the species is unified as I_inactive_ = {j}_j≠i_.

5. A regression for y_i_ by {x_k_}_k∈I_active__ is performed by S-map. This returns *MSE_prev_*.

6. For each index j in I_inactive_, create 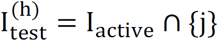, where the suffix h indicates that j is the hth element of I_inactive_.

7. Perform a regression y_i_ by 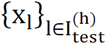 by S-map.

8. Repeat 7 to obtain *MSE*^(*h*)^ for all h.

9. Set the least *MSE*^(*h*)^ as *MSE*_*best*_ and 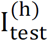 as I_best_.

10. If Q = 1 − *MSE_prev_*/*MSE_best_* is greater than a pre-specified value (Q_c_) then set *MSE_best_* as *MSE_prev_* and I_best_ as I_active_, remove hth element of I_inactive_ and go back to 6, otherwise go to 11.

11. Return 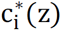 where z indicates that the inferred interaction strength for species i is obtained by the zth iteration. If z < Z increment z by 1 and go back to 3, otherwise terminate the loop.

12. Return 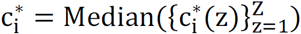.

Q_c_ controls the sensitivity of the algorithm to find links between species. It is reasonable to fix Q_c_ as zero to include a new link when it improves predictability of y_i_s. However, very small improvement of Q_c_ would result in inclusion of erroneous links. For this reason, we set Q_c_ as 0.01 in both the SSM and LIMITS.

### Simulation model

We used a population dynamics model to generate the data set for validation. The model is based on a generalized Lotka-Volterra equation (GLVE), 
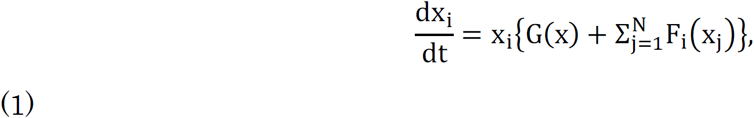
which has been frequently used to model microbial population dynamics (Fisher and Mehta 2014; Coyte et al. 2015; Bucci et al. 2016). We assumed that 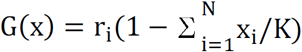, where r_i_ is the intrinsic growth rate and K is the carrying capacity that defines upper limit of abundance. For F_i_, to show that our approach actually is equation-independent, we tested three different forms, 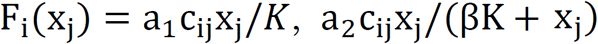 and 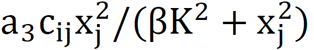, known as Holling type I, II and III functional response, respectively (see supplementary information). In F_i_, c_ij_ controls the per capita effect of species j on i. Hence, the matrix C = {c_ij_} represents the presence or absence of interaction between species. β is the half-saturation constant of the interspecific interaction.

To reduce dimensionality of the right hand side of equation (1), we used the relationship,

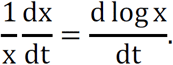
Hence eq. (1) is transformed to, 
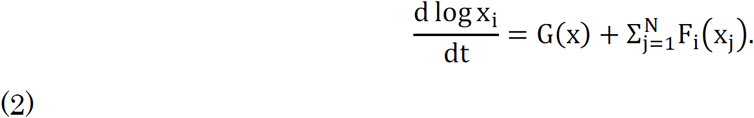

Then, with a noise term, eq. (2) is discretized as,

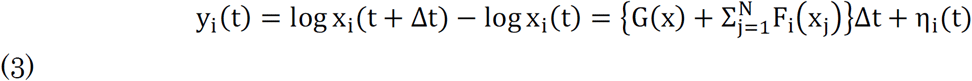

This equation gives the true relationship between x’s and y_i_ when applying the regression test. Here, η_i_(t) is a random vector whose elements are drawn from a normal distribution with mean 0 and standard deviation 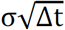. We used three different values (0.01, 0.1, 0.2) for σ to test the effect of the different strength of stochasticity. Moreover, to simulate observational errors, each abundance 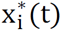 was perturbed by a random value drawn from a normal distribution with mean zero and standard deviation 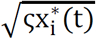. Here, we used four different values (0,1,4,16) for ς.

Please note that due to (3), the inferred interaction strength 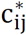 corresponds to the time average of F_i_(x_j_)/x_j_ instead of c_ij_ itself. However, we later confirm that there is a strong correlation between 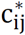 and c_ij_ as a whole. We emphasize that although we specify G and F here, the SSM does not require F and G to be known or even described as a specific mathematical formulation. Please see supplementary information for the details of the data generation procedure.

### Application of the SSM to mouse microbiome data

We applied the SSM to the time-series data of gut-microbiota taken from the faeces of six male C57BL/6J mice (M1 to M6) which were maintained in a vinyl isolator over a 72 week period (Nakanishi Y. et. al, unpublished data). Feaces were sampled once every 4 weeks between 4 to 72 weeks of age, thus 18 data points were obtained per mouse. In the mouse gut microbiome data we used, OTUs were categorized into genus-level groups by the CLASSIFIER program of the Ribosomal Database Project (RDP) within a software package Quantitative Insights into Microbial Ecology (QIIME). For our analysis, we picked the seven most abundant groups that comprise approximately 85% of the total microbial biomass and classified the abundance of the remaining groups into a single group of “Others”. Next, we calculated (x, y_i_) for all mice and selected the dataset of a mouse (M_k_) as *D_tg_*. At each iteration of the bagging procedure, half of the dataset points of the remaining five mice were randomly sampled as *D_db_*. The above procedure was repeated for k=1 to 6.

### Performance criteria

Two major performance criteria for network inference methods are sensitivity (the ratio of detected interacting species pairs with respect to all interacting pairs) and specificity (the ratio of detected non-interacting species pairs with respect to all non-interacting pairs; Fig. S2). Furthermore, accuracy (the ratio of detected interacting species pairs plus that of the non-interacting species pairs with respect to all possible species pairs) quantifies the overall performance of the method for discriminating interacting and non-interacting pairs. These are binary classifications that indicate only the correctness of identifying absence or presence of interaction. We confirmed that these were sufficient because there was a strong correlation between inferred and actual interaction strength.

### Results

We compared the performance of the SSM and a comparative equation-based method (LIMITS) in a five-species coupled food chain model (Fig. 2a, b). The model was introduced by Deyle et al. (2016) to show the ability of S-map to forecast complex population dynamics. In their study, however, the interaction topology was assumed to be known. Our results showed that even without prior knowledge of the interaction topology, the SSM was able to recover the actual topology from time-series data if the dataset was large enough. Figure 2c shows that the accuracy of SSM outperformed that of LIMITS when the dataset was larger than 50 points. The increase in dataset size improved sensitivity of both methods (Fig. 2d), while it reduced the specificity of LIMITS (Fig. 2e). Hence, the SSM was better at avoiding false positives.

**Figure 2.**
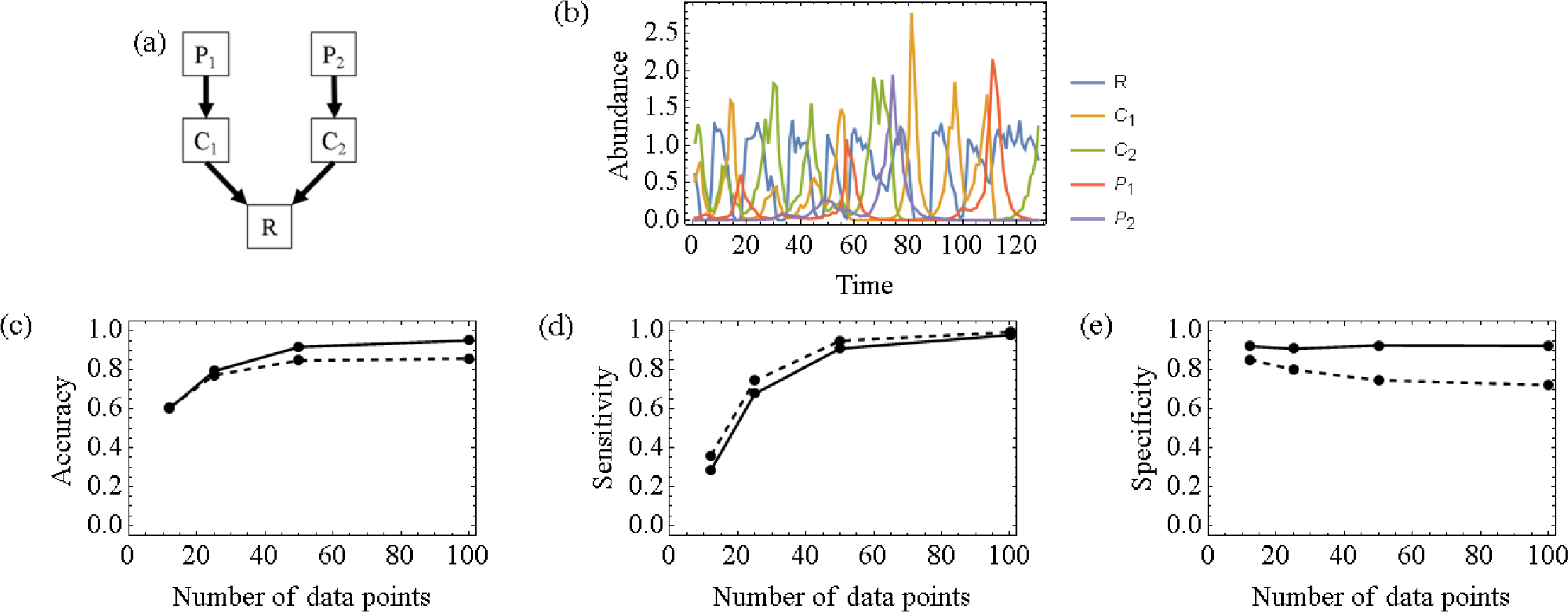
Results of a five species coupled food chain model. Interaction topology of the model (a) and population dynamics (b). Accuracy (c), sensitivity (d) and specificity (e) of the SSM (solid) and LIMITS (dashed) as a function of the number of data points was calculated by assembling 100 different trials. Please see supplementary information for the details of the five species model and simulation.

To compare the SSM and LIMITS further, we tested their performance for complex ecological dynamics generated from a generalized Lotka-Volterra model with seven species with random interactions. A significant positive correlation (p < 0.001) between inferred interaction strength (cij) and that of the actual network (c_ij_) was found in both the SSM and LIMITS, although the SSM showed a stronger correlation than LIMITS (Fig. 3). Figure 4 shows that increased dataset size improved the accuracy of the SSM, while that of LIMITS remained almost constant. These results were robust for the different types of functional responses. The performance of the SSM was equivalent to that of LIMITS even in the worst case. Interestingly, the increase in the magnitude of process noise improved accuracy both in the SSM and LIMITS when the inter-specific interaction was a type II or III functional response, while it reduced accuracy both in the SSM and LIMITS when the inter-specific interaction was a type I functional response. Next, we tested the effect of observational errors by setting the process noise as σ = 0.2 (Fig. 5). An increase in dataset size improved accuracy only in the SSM, as shown in figure 4. Similarly, a decrease in the magnitude of observational errors improved the accuracy of the SSM, while that of LIMITS remained almost constant over all cases. Here again, the performance of the SSM was equivalent to LIMITS even in the worst case. In all cases above, the SSM had greater specificity than SLR, which compensated for its lower sensitivity (Fig. S3-6).

**Figure 3.**
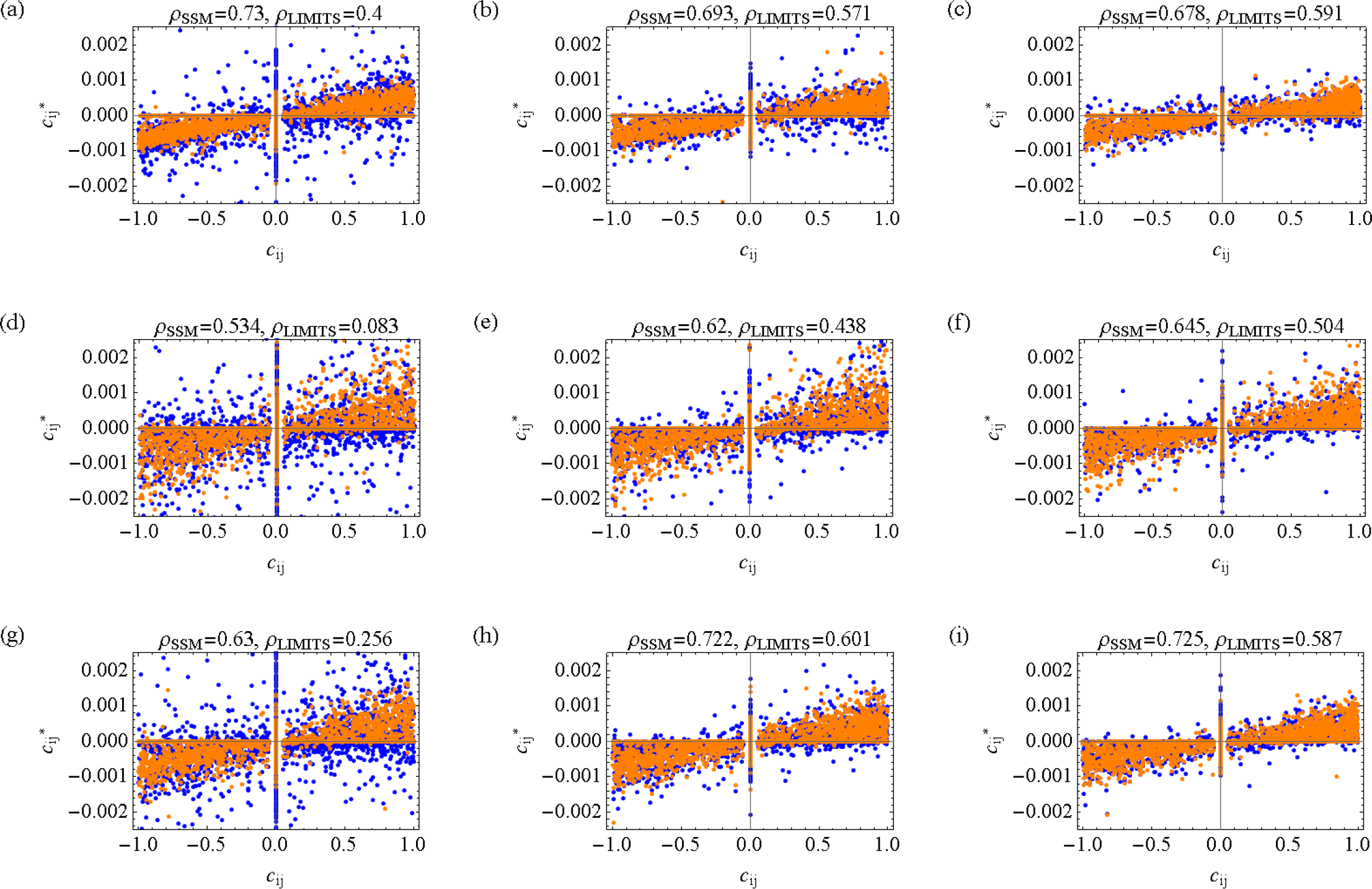
Scatter plot of 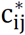 and c_ij_, and correlation measured by Pearson’s correlation coefficient (orange: SSM, blue: LIMITS). (a-c) Holling type I functional response, (d-f) Holling type II functional response, and (g-i) Holling type III functional response with process noise 0.01, 0.1 and 0.2. Points were assembled from 100 network inference results calculated from 100 different time-series data with 100 dataset points generated from a seven species GLV model with random species interactions, σ = 0.01 and ζ = 0.

**Figure 4.**
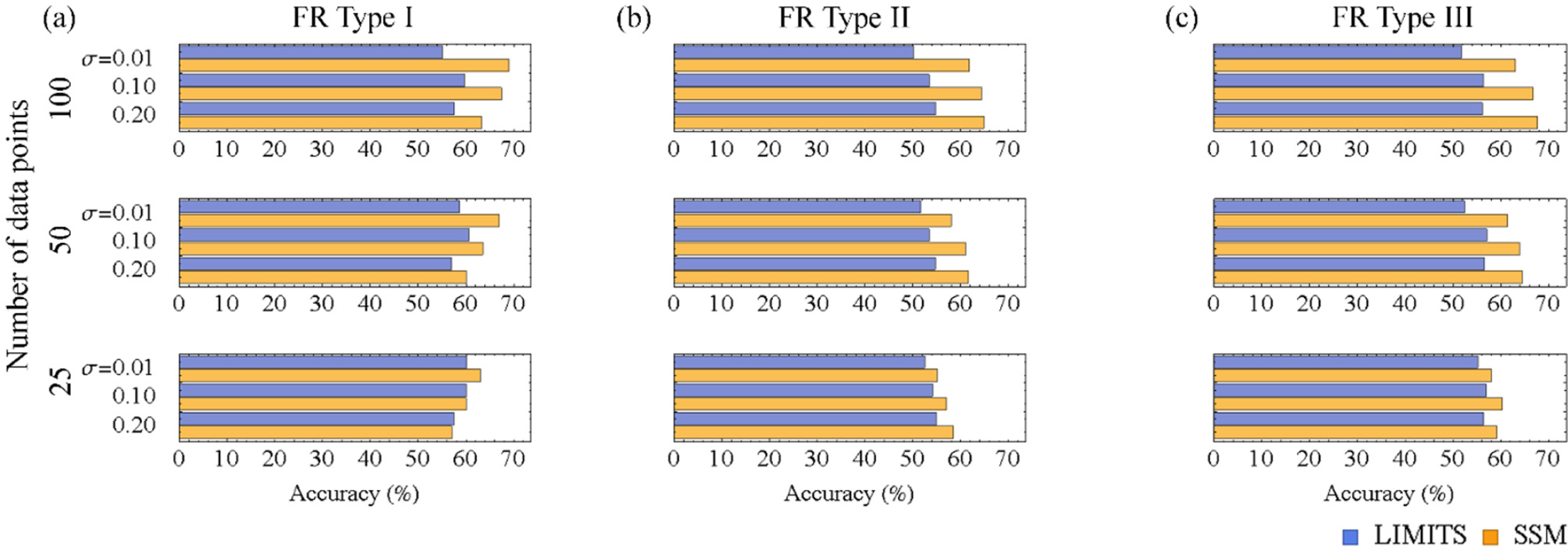
Accuracy of the SSM and LIMITS for the inter-specific interactions of simulated ecological dynamics with Holling type I (a), II (b) and III (c) functional response calculated by assembling 100 different network inference results (calculated from 100 different time-series data with100, 50 and 25 dataset points generated from a seven species GLV model with random species interactions and different magnitudes of process noise, ζ = 0).

**Figure 5.**
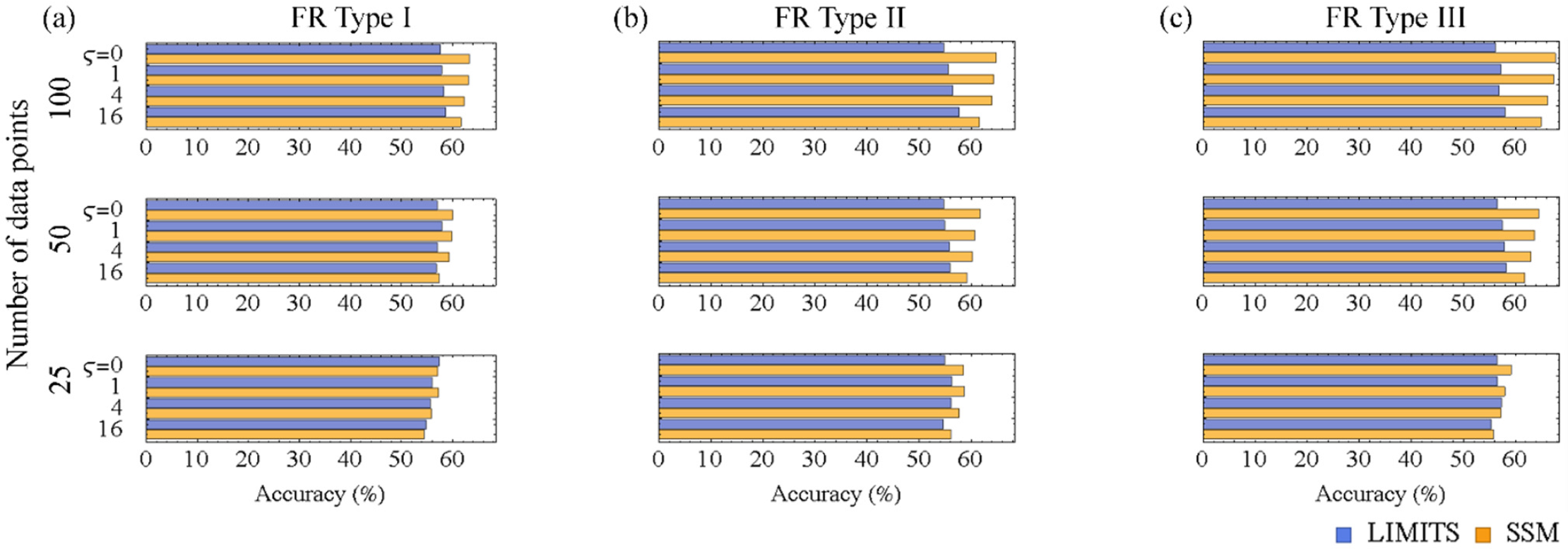
Accuracy of the SSM and LIMITS for the inter-specific interactions of simulated ecological dynamics with Holling type I (a), II (b) and III (c) functional responses calculated by assembling 100 different network inference results (calculated from 100 different time-series with100, 50 and 25 dataset points generated from a seven species GLV model with random species interactions and σ = 0.2 perturbed by different magnitudes of observational errors).

The ecological interaction networks of mouse gut microbiota were inferred using the complete dataset from weeks 4-72 as well as comparing weeks 4-40 and weeks 36-72 to consider the shift in community composition around the middle of the mouse’s aging processes (Langille et al. 2014; Nakanishi Y. et al, unpublished data). The number of possible links identified by the SSM was 40 for time-series data from weeks 4-72, 19 for weeks 4-40 and 16 for weeks 36-72, including overlap among different mice. The number of identified links in weeks 4-40 and weeks 36-72 week was expected to be smaller than that of weeks 4-72 due to the reduced number of data points. In the network inferred by the data points at 4-72 weeks of age (Fig. 6a), unclassified Rikenellaceae, Ruminococcaceae and Clostridiales had more interaction links than others in this order. These groups varied according to in-degree (effect from other species) and out-degree (effect on other species). The in-degree exceeded out-degree for Rikenellaceae, out-degree was zero while in-degree was 12 for Clostridiales, and both in-degree and out-degree were the same for Ruminococcaceae. The number of positive and negative links was mostly the same in the three species for both in- and out-degree except for Rikenellaceae with 12 negative and 2 positive effects on other species. The comparison of networks inferred by data points at 4-40 weeks of age (Fig. 6b) and 36-72 weeks of age (Fig. 6c) showed that links with Ruminococcaceae were most common in weeks 4-40 while links with Rikenellaceae were most common in weeks 36-72. Clostridiales had positive effects from Ruminococcaceae in weeks 4-40 and negative effect from Rikenellaceae in weeks 36-72. Hence, most of the links with Clostridiales were explained by links from these two species. *Allobaculum* was also a major counterpart of Ruminococcaceae at 4-40 weeks of age.

**Figure 6.**
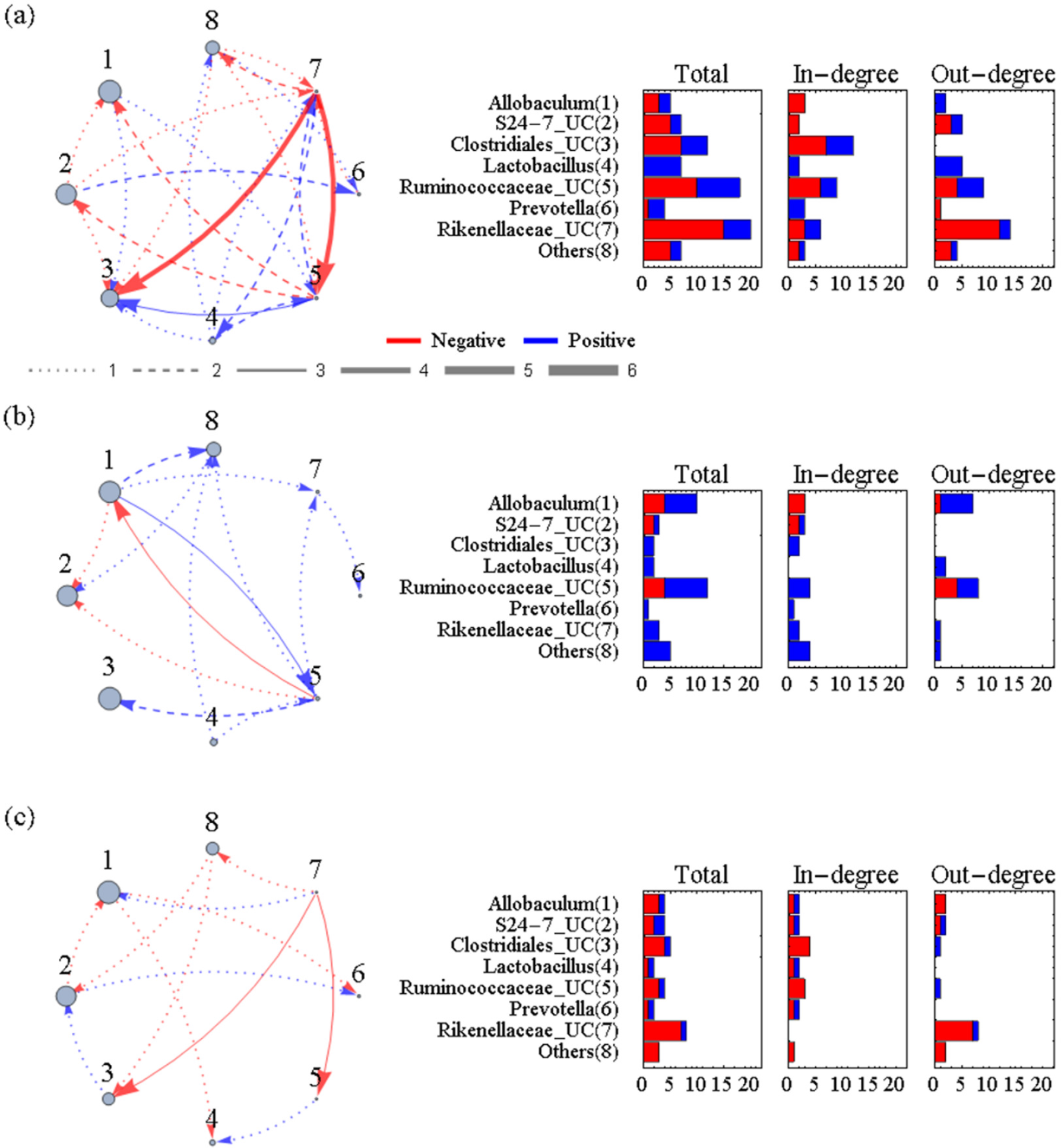
Union of the inter-specific interaction networks of six mice inferred by data points at 4-72 weeks of age (a), 4-40 weeks of age and 36-72 weeks of age (c). The size of the nodes indicates the relative abundance averaged over six mice. UC stands for “unclassified” which means that the OTU was not classified into a known genus.

## Discussion

We developed the sparse S-map method (SSM), an equation-free method used for inferring ecological interaction networks from a multivariate ecological time-series (Fig. 1). Using simulated multispecies population dynamics, we compared the performance of the SSM to a comparable equation-based method (LIMITS), to highlight the differences between equation-free and equation-based methods when applied to complex microbial dynamics (Fig. 2,3,4 and 5). We show that either an increase in dataset size or a decrease in observational error improved the accuracy of SSM (Fig. 4 and 5). On the other hand, the accuracy of LIMITS was almost unchanged for both cases and equivalent to the SSM at best. Hence, the SSM outperformed LIMITS when datasets were large or the magnitudes of observational errors were small. The results were robust to the magnitudes of process noise and functional forms of inter-specific interactions that we tested. These results suggest that the SSM is able to extract more information from time-series data than LIMITS. This is partly due to the equation-free property of the SSM that allows inclusion of detailed descriptions of species relationships into the network inference. However, the SSM had better performance than LIMITS even for a Holling type I functional response that has no explicit non-linearity in its functional form (see supplementary information). In this case, the advantage of the SSM would be explained by its feasibility for the non-equilibrium population dynamics. It is worth discussing how well our approach scaled up to communities with a larger number of species. In principle, applicability of our method is not limited by the number of species. However, the application to a community with a large number of species would require many data points to cover its possible states. Our results suggest that shortage in dataset size reduces sensitivity rather than specificity (Fig. S3-6). Hence, the model will be able to identify some actual interactions even if the number of species is very large compared to the available dataset size.

We applied the SSM to a time-series of gut-microbiota taken from the faeces of six mice. Our results are summarized as follows (Fig. S7): an unclassified Ruminococcaceae commonly affected other species from young to middle age, whereas Rikenellaceae commonly affected other species from middle to old age. Moreover, the Rikenellaceae itself had a negative effect on the Ruminococcaceae from middle to old age. An unclassified Clostridiales was affected by both groups throughout young to old age, while *Allobaculm* was a major counterpart of the Ruminococcaceae from young to middle age. The above mentioned relationship was still found in the network inferred by LIMITS, although it was obscured by other interactions (Fig. S8). These observations reflect the backdrop of lifelong dynamics of gut microbiota. For example, Rikkenellaceae is reported as a common group in middle- to old-aged mice (Langille et al. 2014). Hence, although the unclassified Rikenellaceae was not a dominant group, as a member of this family, it makes sense that it actively interacts with other species in the latter half of the life-stage. Moreover, it is interesting to mention that a high-fat diet decreases the proportion of Ruminococcaceae and increases that of Rikenellaceae (Daniel et al. 2014). The authors explained this based on the role of Ruminococcaceae as a major user of plant polysaccharides. Hence, our results might represent the age related shift in microbial interactions, and possibly its function, relative to nutrient metabolism. It is unclear whether differences in the interaction networks among mice were due to the nature of microbial interaction in the mouse gut microbiota, or simply due to data limitations. The universality of the interaction networks of mouse gut microbiota (Bashan et al. 2016) and the inherent dynamics must be answered in future studies. Recently, Odamaki et al. (2016) showed the age related compositional shifts in human gut microbiota. We anticipate that applying SSM to human subjects in different age groups will offer deeper insights into how the human gut microbiota is shaped through its lifelong developmental processes. For example, these analyses could reveal shifts in relationships between essential functional groups as well as their relationships to the development and ageing of physiological and immunological functions of our body.

While we used only difference of functional responses to control the nonlinearity of species relationships (see supplementary information), a variety of processes will be the source of non-linear species relationships in empirical microbial communities. The complex interdependency of metabolism (Baran et al. 2015), variety in strategies for inter specific competition (Hibbing et al. 2010; Ghoul and Mitri 2016), intercellular signaling such as quorum sensing (Atkinson and Williams 2009), formation of multi-species complexes known as biofilms (Stoodley et al. 2002), and evolutionary processes running concurrently to ecological processes (Gomez et al. 2016; Toju et al. 2017) might all contribute to the mechanistic basis. The relative performance of the SSM to LIMITS will depend on the ubiquity and strength of these processes. In this case, large non-linearity indices characterized the dynamics of mouse gut-microbiota (Fig. S9), indicating the effects of nonlinear relationships. Together with the performance of the SSM shown here, the need of equation-free modelling approaches for the analysis of microbial dynamics is demonstrated.

In the near future, advances in metagenomic technology will further reduce the cost to collect time-series data and its output will be much more accurate and precise. One important question to ask is whether this will allow the replacement of equation-free approaches with equation-based approaches that utilize advanced modelling techniques (e.g., Brunton et al. 2016). There are two reasons why this seems improbable. First, the complex nature of microbial interactions we have described, even with such data, still present difficult challenges in formulating all the present relationships into mathematical formulations (Hartig and Dormann 2013; Perretti et al. 2013a, b; De Angelis and Yurek 2015). Moreover, time-dependency of species relationships also made it difficult to describe dynamics by a simple mathematical formulation even with longer and more precise time series. Second, a theoretical study proved that finding a precise dynamical equation for a time-series is, in general, computationally intractable even with any amount/quality of data (Cubitt et al. 2012). Conversely, these data advances would simply benefit our approach by promoting its ability to find links between species. Thus, the future development of metagenome technologies would reinforce both the applicability and reliability of equation-free approaches and help improve our mechanistic understanding of microbial communities. We agree with DeAngelis and Yurek (2015), who stressed the value of equation-free modeling approaches for the analysis of complex dynamical systems.

## Data accessibility

The mouse gut microbiome data is available in the DDBJ database (http://getentry.ddbj.nig.ac.jp/) under accession number DRA004786. We used Mathematica 10.2 to implement the SSM and LIMITS, generate simulation data, process mouse gut microbiome data and to perform the analysis. Computer codes (Mathematica notebook files) are available as online supporting information.

## Supplementary information

### Functional responses

Holling Type I functional response is based on the assumption that the handling time required for interaction is negligible, thus F_i_(x_j_) linearly increase with x_j_. On the other hand, type II functional response includes the handling time for interaction and is characterized by a saturation of F_i_(x_j_) when x_j_ increases. Type III functional response considers, as well as the effect of the handling time, a mechanism that suppresses increase of F_i_(x_j_) when j is rare. For example, it may be applicable when a species changes interaction mechanism depending on the relative abundance of other species. Actually, type II functional response is a special case where γ = 1 in 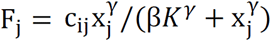 and type III response includes all the cases where γ > 1. Hence, γ is a parameter that controls the nonlinear dependency of F_i_ on x_j_. It is worth noting that in the type II response, β also controls the strength of non-linearity because F_j_/x_j_ = c_ij_/(β*K* + x_j_) is close to c_ij_/β*K* if β*K* is larger than x_j_. On the other hand, type I functional response has no explicit nonlinearity. Hence, in terms of the degree of nonlinearity implemented in the functional form, type III response is the largest followed by type II and type I responses.

### Data generation

Based on eq. (3), we generated ground truth data by numerically solving,

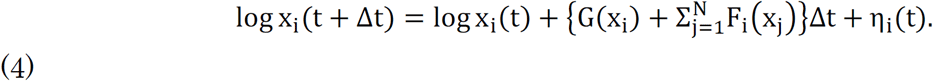

We generated the initial state as 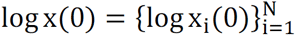 where log x_i_(0) is a random value drawn from a uniform distribution (5,9). The interaction matrix C is generated so that it has M non-zero off-diagonal elements, where value of the non-zero elements is assigned from a uniform distribution (−1, −0.05) or (0.05,1) randomly. Here, we set Δt = 0.1, N = 7 and M = 20. The diagonal elements of C is set to -0.5. With this initial state and interaction matrix, we numerically solved eq.(4) up to 5000 steps and took the latter 1000, 500 or 250 steps. The numerical simulation was discarded if the abundance of at least one species fell below one; otherwise the result was sampled every 10 steps to make a time-series with 100, 50 or 25 data points.

Other parameter values were as follows. We set r = 1 so that the scale of dynamics was relevant to the simulation of microbial dynamics observed as the time-series of 100, 50 and 25 data points mentioned above. K was set to 104 considering the standard size of a metagenomic read count. For, interspecific interactions, we set a_l_ = 7.5, a_2_ = 2.7, a_3_ = 2 and β = 0.1.

To prepare non-equilibrium time-series data, we accepted only a time-series whose minimum value of θ calculated for all species was larger than 1. In this pre-evaluation, we calculate θ by using all species as explanatory variables

### Five species coupled food chain model

The equations of five species coupled food chain model are, 
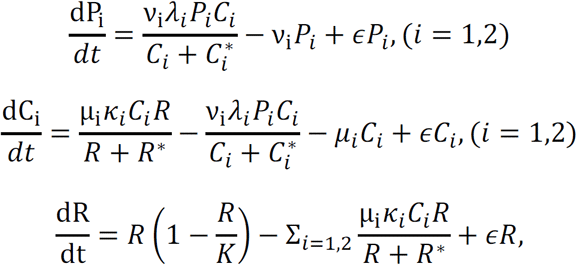
where P_l_ and P_2_ are predator abundances, C_l_ and C_2_ are consumer abundances, and R is the shared resource abundance. Parameter values were set as follows; ν_1_ = 0.1, ν_2_ = 0.07, λ_l_ = 3.2, λ_2_ = 2.9, 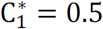, 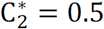, μ_l_ = 0.15, μ_2_ = 0.15, κ_l_ = 2.5, κ_2_ = 2.0, R* = 0.3, k = 1.2. After the transformation as in eq. (1) to (4), we numerically solved the equations with step size Δt = 0.1 upto 5,000 steps and took the latter 1000, 500, 250 or 120 steps. The result was sampled every 10 steps to make a time-series with 100, 50, 25 or 12 data points.

---

**Figure S1.**
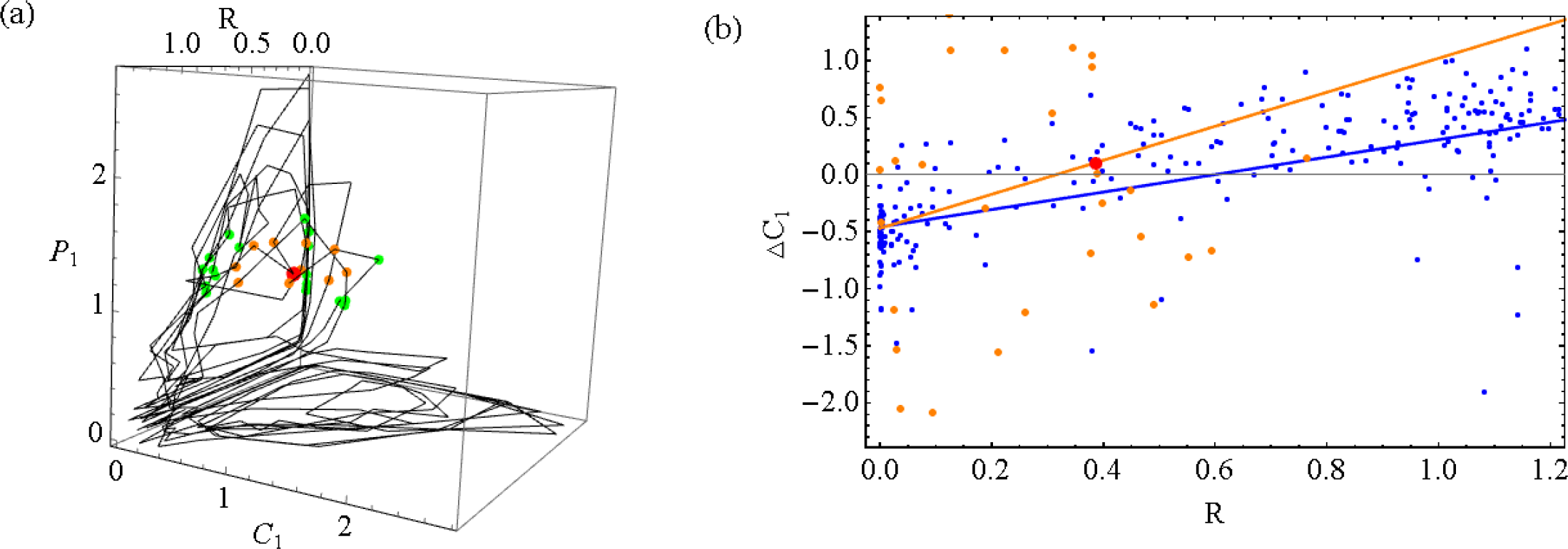
A schematic description of the SSM (S-map). (a) top 10 closest points (orange) and top 11 to 20 closest points (green) of a target state (red) are plotted in a three dimensional space. (b) regression lines in a two dimensional space (R, ΔC_1_) by S-map using R and C_1_ as explanatory variables and ΔC_1_ = logC_1_(*t* + 1) − logC_1_(*t*) as an response variable is shown for θ = 0 (blue) and θ = 1 (orange). Here, top 30 closest points of a target state (red) is also indicated by orange.

**Figure S2.**
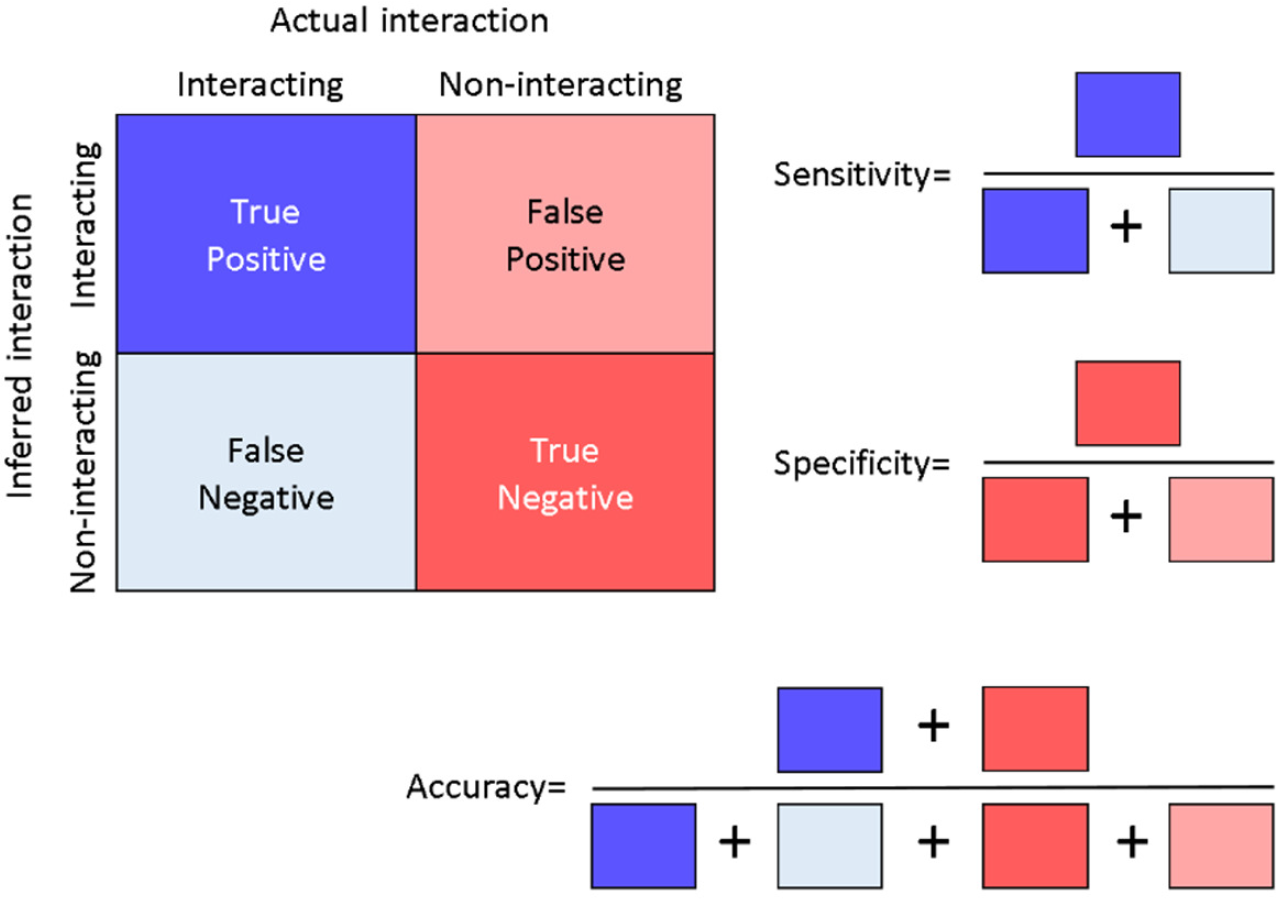
Binary criteria used for evaluation.

**Figure S3.**
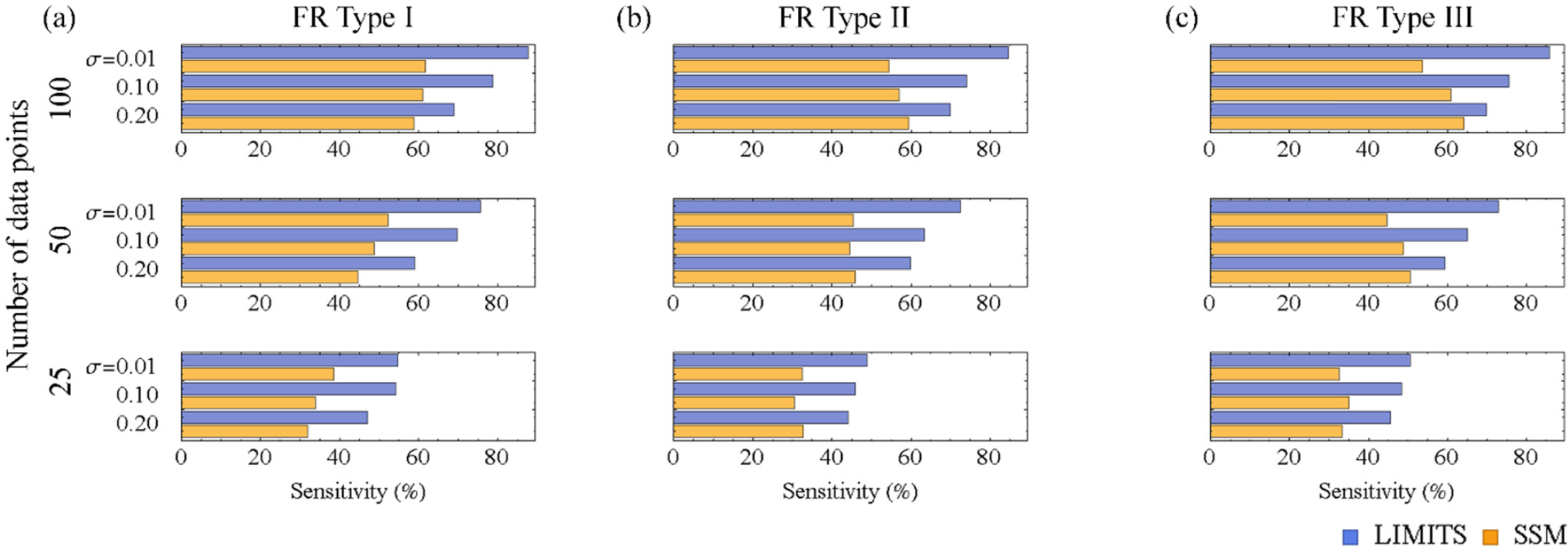
Sensitivity of the SSM and LIMITS for the inter-specific interactions of simulated ecological dynamics with Holling type I (a), II (b) and III (c) functional responses calculated by assembling 100 different network inference results (calculated from 100 different time-series with100, 50 and 25 dataset points generated from a seven species GLV model with random species interactions and different magnitudes of process noise).

**Figure S4.**
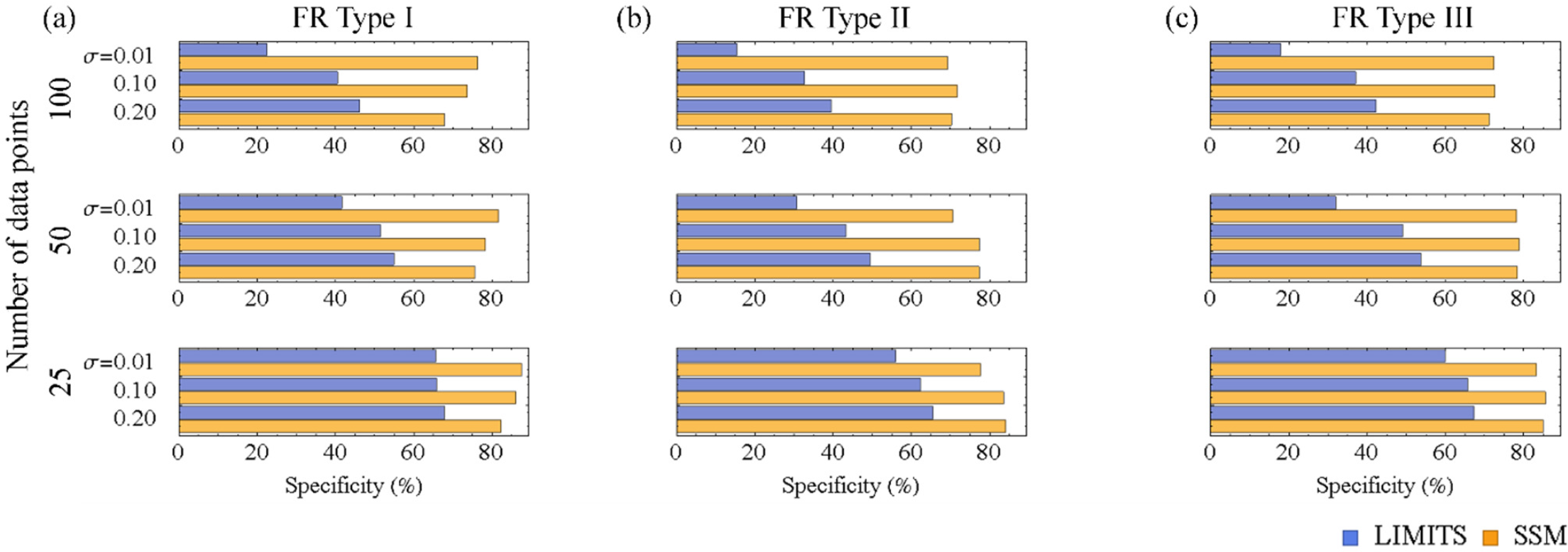
Specificity of the SSM and LIMITS for the inter-specific interactions of simulated ecological dynamics with Holling type I (a), II (b) and III (c) functional responses calculated by assembling 100 different network inference results for different magnitudes of process noise and dataset size (calculated from 100 different time-series with100, 50 and 25 dataset points generated from a seven species GLV model with random species interactions and different magnitudes of process noise).

**Figure S5.**
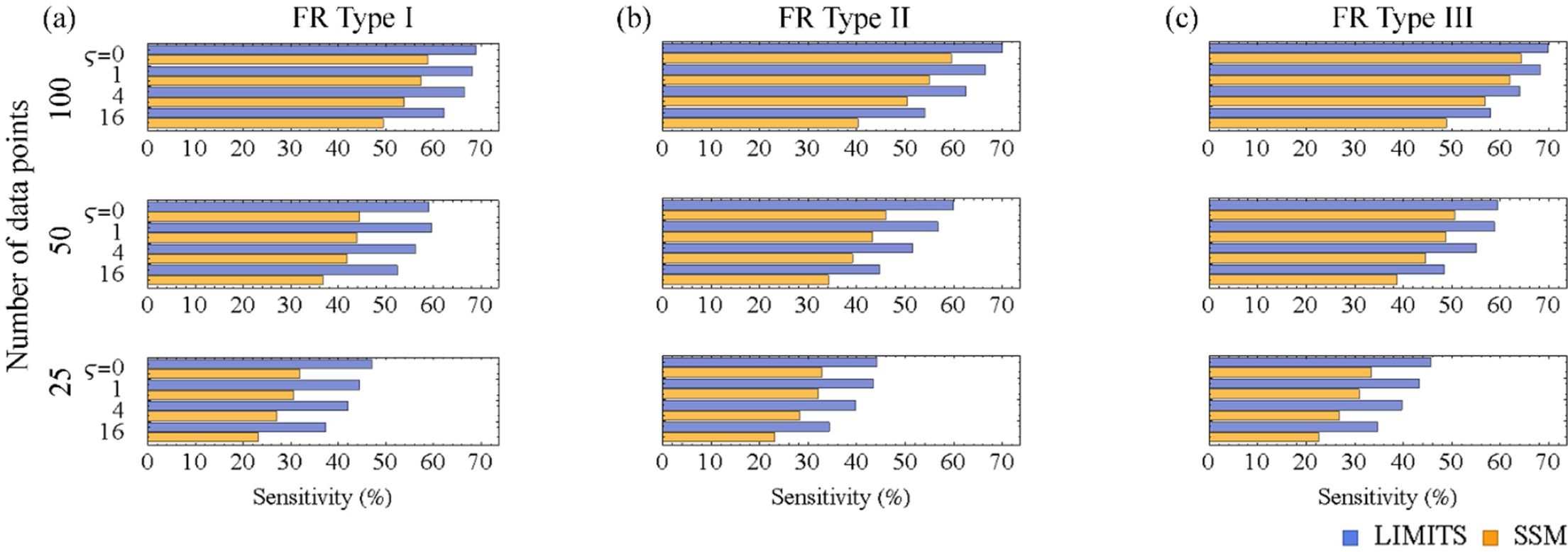
Sensitivity of the SSM and LIMITS for the inter-specific interactions of simulated ecological dynamics with Holling type I (a), II (b) and III (c) functional responses calculated by assembling 100 different network inference results (calculated from 100 different time-series with100, 50 and 25 dataset points generated from a seven species GLV model with random species interactions perturbed by different magnitudes of observational errors).

**Figure S6.**
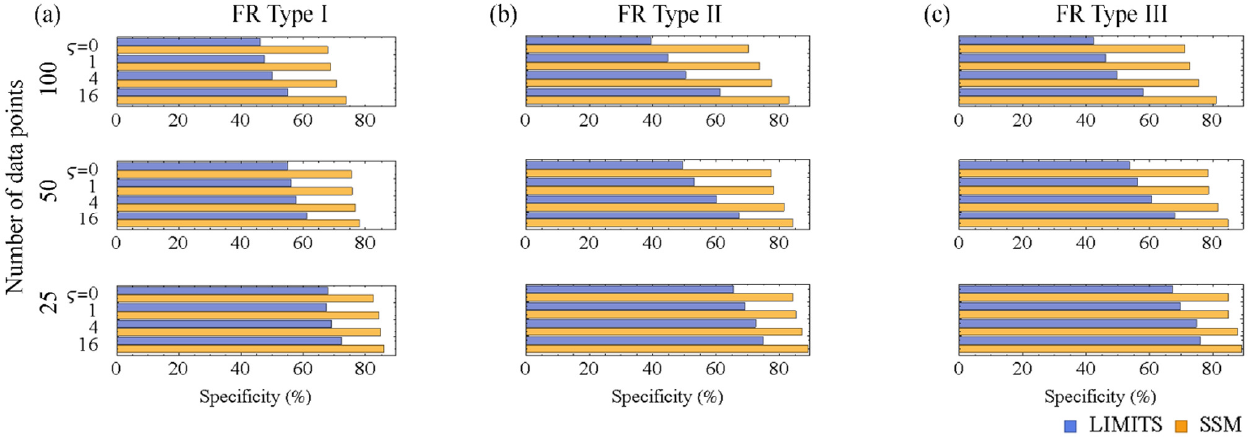
Sensitivity of the SSM and LIMITS for the inter-specific interactions of simulated ecological dynamics with Holling type I (a), II (b) and III (c) functional responses calculated by assembling 100 different network inference results (calculated from 100 different time-series with100, 50 and 25 dataset points generated from a seven species GLV model with random species interactions perturbed by different magnitudes of observational errors).

**Figure S8.**
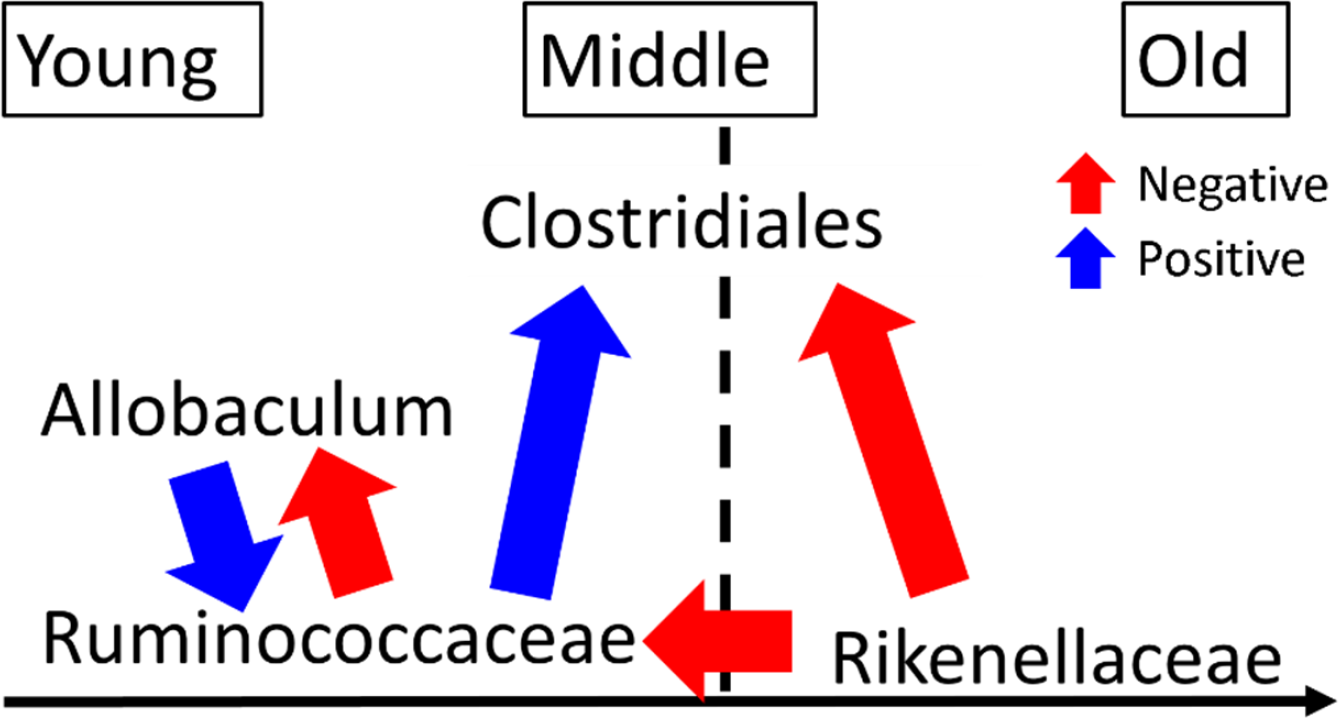
Schematic description of microbial interaction identified by the SSM.

**Figure S7.**
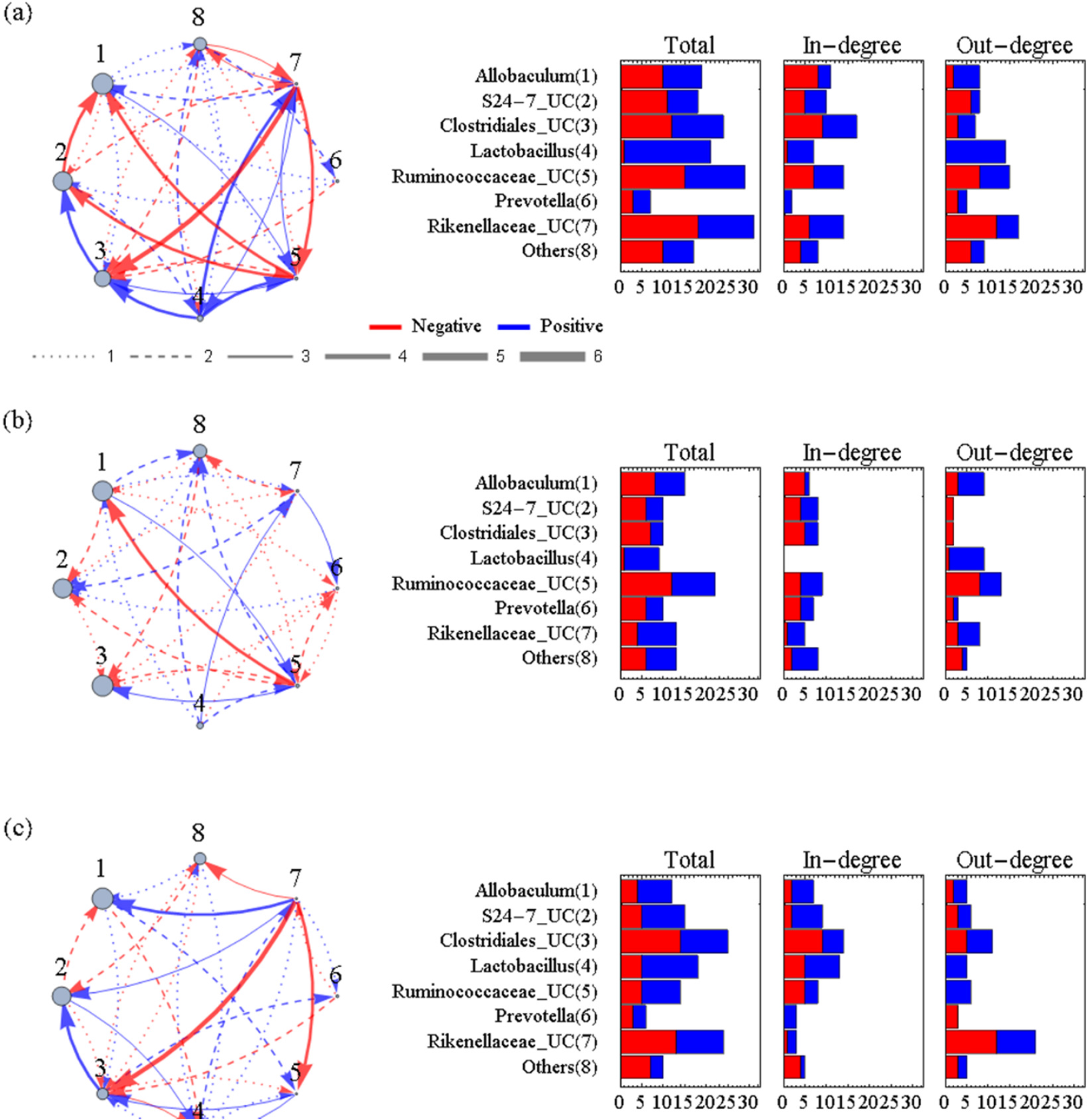
Union of the inter-specific interaction networks of six mice inferred at 4-40 weeks old (a-f). Positive and negative effects are indicated by blue and red arrows respectively. The size of nodes indicates relative abundance. Inter-specific links are excluded. UC stands for “unclassified” which means that the OTU was not classified into a known genus.

**Figure S9.**
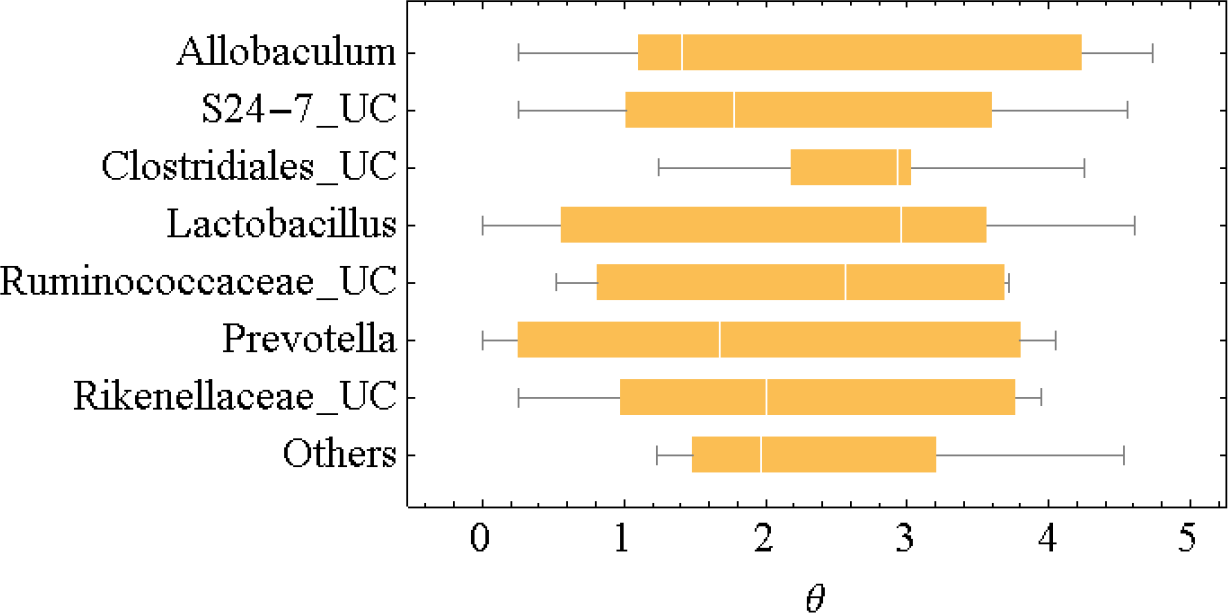
Box plot of the θ of the seven most abundant groups in the mouse gut microbiome for M_1_. θs were not significantly different for other mice.

